# So similar, yet so different: the paradigm of PARP9 macro domain paralogs

**DOI:** 10.64898/2026.08.12.744356

**Authors:** Nikolaos K. Fourkiotis, Christos Sideras-Bisdekis, Aikaterini C. Tsika, Alexander Fish, Konstantina P. Kravvariti, Sofia-Antigoni Tsatsouli, Anastassis Perrakis, Aleksandra Chikunova, Georgios A. Spyroulias

## Abstract

Human PARP9 harbours two tandem macro domains, MD1 and MD2, with distinct roles in ADP-ribosylation signaling. Whereas MD1 is a MacroD-type de-MARylase (“eraser”), MD2 functions as a MacroH2A-like “reader” lacking detectable hydrolase activity. How two domains so similar in sequence and fold achieve such divergent functionality has remained unclear. Our de-MARylation assays confirmed this division of labor, even though crystal structures revealed nearly identical α/β/α folds and similar binding pockets, and solution NMR shows broadly comparable dynamics. A remarkable distinction, however, emerges from isothermal titration calorimetry, showing that both domains bind free ADP-ribose with comparable affinity (K_D_ = 5.4 and 8.4 μM), but through markedly different thermodynamics: MD1 binding is enthalpy-driven and offset by a larger entropic penalty, whereas MD2 binds with weaker enthalpy and a smaller entropic cost. Our ADPr-bound crystal structures rationalized this showing that the distal ribose is positioned differently in the two pockets, connecting to a glycine-rich catalytic loop and a conserved aromatic residue present in active macro domains like MD1, but altered in MD2, while the catalytic asparagine itself is structurally conserved (Asn140/Asn339). Moreover, we show that a single amino acid substitution in MD2 leads to detectable RNA de-MARylation activity without altering its fold. Together, our findings show that an eraser-versus-reader distinction between the macrodomains in PARP9 is encoded in the dynamics of ligand binding and the exact position the distal ribose, revealing an unexpected catalytic plasticity relevant to PARP9’s roles in immunity and cancer.

## Introduction

Macro domains (MDs) are evolutionarily conserved protein modules that mediate interactions with ADP-ribose (ADPr) and related NAD^+^ metabolites. They typically contain a positively charged binding pocket and adopt a conserved α/β/α sandwich fold. Initially referred to as “X-domains” due to their unknown function, their structural similarity to the histone variant MacroH2A later led to the designation “macro domains”^1^. MDs are widely distributed across all domains of life, including bacteria, archaea, viruses, and eukaryotes.

Functionally, macro domains are key regulators of ADP-ribosylation, a post-translational modification (PTM) involved in essential cellular processes such as transcriptional regulation, DNA repair, and immune signaling^2^. MDs can act as “readers” selectively recognizing free ADP-ribose (ADPr) or ADP-ribosylated substrates, and/or as “erasers” catalyzing the hydrolysis of covalently attached ADPr moieties. Some domains may fulfill both roles. MDs are classified into six types^3^, of which four major types are found in humans: MacroH2A-like, ALC1-like, MacroD-type, and PARG-like. In contrast, viruses and some yeasts encode additional MD types, including SUD-M-like and Macro2-type^4^. In humans, macro domains are present in at least eleven proteins, including CHD1L, GDAP2, MACROH2A1, MACROH2A2, MACROD1, MACROD2, TARG1, PARG, PARP9, PARP14, and PARP15. Among human macro domain-containing proteins, PARP9 is distinctive in that it harbors two tandem macro domains (MD1 and MD2) but lacks ADP-ribosyltransferase (ART) activity due to substitutions in the catalytic triad of its ART domain^5^. ART activity is the catalytic transfer of ADPr from NAD⁺ onto substrate side chains (amino acids, or nucleic acid termini), attaching single units (mono-ADP-ribosylation-MARylation) or chains (poly-ADP-ribosylation-PARylation) of ADPr. PARP9 belongs to the macro-PARP subfamily together with PARP14 and PARP15, which are characterized by the presence of multiple macro domains N-terminal to the ART domain^6^.

PARP9 serves as a noncanonical sensor of RNA viruses by recognizing long double-stranded RNA (dsRNA, 1100-1400 bp), thereby promoting type I interferon (IFN) production^7^. This mechanism likely extends to single-stranded RNA (ssRNA) viruses like SARS-CoV-2, which produce dsRNA intermediates during replication. PARP9 activation triggers innate immune pathways, even in the absence of canonical signaling through mitochondrial antiviral-signaling protein (MAVS)^7^. PARP9 is also involved in targeting viral proteins for ubiquitination, thus amplifying antiviral gene expression^8^. Beyond antiviral immunity, PARP9 contributes to host defense against bacterial infections. During *Mycobacterium tuberculosis* infection, PARP9 expression is upregulated, and it suppresses the intracellular bacterial growth. Reintroduction of PARP9 into deficient lung epithelial cells reduces infection efficiency and bacterial proliferation, potentially through modulation of cytokine responses and NF-κB signaling^9^. In addition, PARP9 has been implicated in cancer progression. It is overexpressed in aggressive lymphomas such as diffuse large B-cell lymphoma (DLBCL), where it contributes positively to tumor survival and immune evasion ^10^. In breast cancer, elevated PARP9 levels (observed in ∼44% of cases) are associated with increased cell migration, suggesting a role in metastasis. Similarly, in gliomas, PARP9 overexpression correlates with higher tumor grade and poorer prognosis, indicating its potential as a prognostic marker and therapeutic target^11^.

PARP9 forms a stable complex with the E3 ubiquitin ligase DTX3L^8^ and is rapidly recruited to sites of DNA damage in a manner dependent on ADP-ribosylation by PARP1 and PARP2^10,12^. Early studies found MD2 to be essential for PARP9 localization to DNA damage sites^10^, while later work showed that this recruitment is disrupted also when MD1 is mutated^12^. These observations underscore the functional importance of the tandem macro domain architecture in PARP9, yet the molecular basis for their distinct ligand-binding and catalytic roles has remained unclear. The two PARP9 macro domains exhibit distinct functional properties. MD1 is a MacroD-type domain with de-MARylation activity, capable of hydrolyzing mono-ADP-ribose from ADP-ribosylated substrates such as proteins and nucleic acids^13^. In contrast, MD2 is a MacroH2A-like domain that lacks detectable hydrolase activity^13^, functioning instead as a high-affinity ADPr-binding module. These findings suggest a division of labor in which MD1 acts as an eraser of ADP-ribosylation while MD2 functions as a reader. How these two domains cooperate within a single protein, however, remains poorly understood.

To fully understand PARP9 MD1 and MD2 functionality and their functional cooperation, information about their structures and interactions with ligands is indispensable. However, structural insight into PARP9 macro domains has been limited, with only the apo structure of MD2 being previously available. To address this gap, we applied an integrated structural biology approach combining X-ray crystallography, NMR spectroscopy, biophysics, and biochemical assays to investigate the structure-function relationship of both PARP9 macro domains.

Biophysical and biochemical studies confirmed that only MD1 exhibits de-MARylation activity towards auto-MARylated PARP10 ART domain and ADP-ribosylated RNA. The crystal structures of MD1 and MD2 in their apo and ADPr-bound forms reveal marked differences in ligand binding and positioning inside the binding pocket. Consistent with their structural features, structure driven site-directed mutagenesis of the binding pocket residues, in both domains, highlighted their role in modulation of ligand binding and catalytic efficiency. Collectively, these findings enhance our knowledge of PARP9’s function in modulating ADP-ribosylation via its two MDs. We offer new insights towards understanding the functional significance of PARP9 in immunity and cancer biology.

## Results and Discussion

### MD1 is an active mono-ADPr hydrolase, while MD2 is catalytically inert

To confirm that the isolated MD1 domain functions as an eraser and MD2 acts as a reader of ADP-ribosylation ^12–14^ we performed activity assays measuring removal of ADPr from auto-MARylated PARP10 (Fig. 1a). MD1 removed ADPr at an apparent rate of 0.130 ± 0.003 μM/min, whereas MD2 showed no detectable activity over 60 min. Addition of free ADPr as a competitive inhibitor (1:250) slows down MD1 to 0.078 ± 0.009 μM/min (Supplementary Fig. 1), confirming that the substrate and free ADPr compete for the same pocket. In the physiologically relevant tandem MD1-MD2 construct, the rate of hydrolysis by MD1 modestly decreased to 0.110 ± 0.004 μM/min, indicating that MD2 could compete with MD1 for binding.

**Figure 1.**
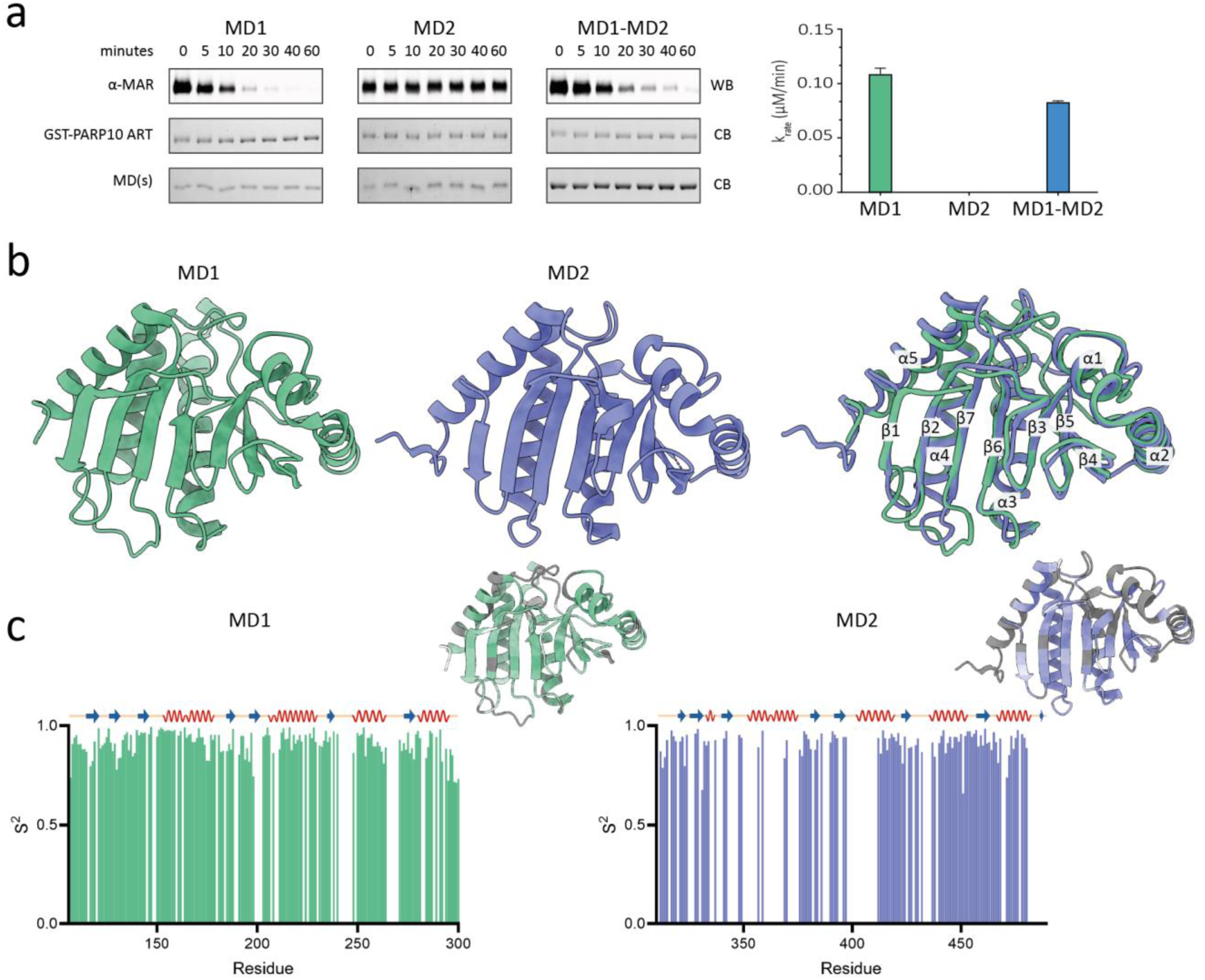
Biochemical and structural characterization of the tandem macro domains of PARP9 MD1 and MD2. **a.** De-MARylation activity of MD1, MD2, and a double domain construct (MD1-MD2) against GST-PARP10 ART, monitored by anti-MAR Western blot and Coomassie staining over 60 minutes (left) with calculated reaction rates (right). Data are mean with SD, n=3. **b.** Crystal structures of MD1 in green (9QYD) and MD2 in purple (9QYG) and their overlay. **c.** Backbone dynamics: Lipari-Szabo order parameters (S^2^) plotted against residue numbers for MD1 (green) and MD2 (purple), with secondary structure elements indicated above and S^2^ values mapped onto the 3D structures (gradient from white to darker green/purple = less rigid to more rigid; grey = unassigned residues.

### The structural investigation of the domains indicated slight but functionally relevant differences

The structures of the MD1 and MD2 macro domains of PARP9 were determined by X-ray crystallography to 1.91Å and 2.7Å resolution respectively (Fig. 1b), providing the first experimental structure of PARP9 MD1. Both adopt the canonical macro domain fold, characterized by a three-layered α/β/α sandwich architecture, with a layer of seven-stranded β-sheet flanked by two and three α-helices on either β-sheet side. In the MD1 structure, electron density was poorly defined for residues G102-K105 at its N-terminus, and at a loop between the secondary elements α4-β7, probably due to local flexibility. In the MD2 structure, regions with missing electron density included residues T305-F309 (N-terminus), K370-S375 (loop between α2-β4), and S491-V497 (C-terminus).

Overall, the secondary structure elements of the two macro domains are highly conserved and aligned. Despite their large functional differences, the two macro domains display only minor structural differences. A notable structural distinction between MD1 and MD2 lies in the third α-helix, located between the fifth and sixth β-strands (β5-α3-β6). In MD1, this helix spans 26 amino acids; six more than the corresponding 20-residue helix in MD2. Tunnel analysis (MOLEonline^15^) revealed that despite comparable overall cavity volume, MD1 (783.1 Å^3^) and MD2 (795.6 Å^3^) differ markedly in entrance geometry: MD1 possesses a narrower channel (1.0 Å bottleneck radius), than MD2 (1.5 Å), potentially facilitating more rapid ligand access consistent with a reader function^16^.

Across macro domains, ADPr recognition and de-MARylation depend on a conserved set of structural elements. The adenosine is clamped by a rigid β2–β3 region, the diphosphate by a basic loop, and the distal ribose, the site of chemistry in active de-MARylases, is accommodated by a flexible, glycine-rich catalytic loop carrying a conserved asparagine that, together with an ordered water, mediate hydrolysis. The catalytic asparagine is present and structurally conserved in both PARP9 macro domains (Asn140 in MD1, Asn339 in MD2), arguing that MD2’s lack of activity does not stem from loss of this catalytic residue. The surrounding loops, however, diverge. In MD1 the catalytic asparagine is followed by an acidic Glu141–Asp142 pair, whereas in MD2 it is followed by Pro340–His341. The glycine-rich loop of MD1 (Gly146-Gly147-Gly148) is replaced in MD2 by Val345-Gly346-Pro347, losing two glycines and introducing two other hydrophobic residues with bulky side-chain and reduced conformational flexibility due to the proline.

To better understand the functional differences, we analyzed the dynamics of two PARP9 macro domains by solution NMR. Backbone dynamics analysis (^15^N R₁, R₂, and {^1^H}-^15^N NOE) revealed similar ps-ns dynamics for apo MD1 and MD2, for the assigned residues, consistent with a compact monomeric protein state in solution (Supplementary Fig. 2). MD1 displayed a rotational correlation time of ∼11.4 ± 0.63 ns with high heteronuclear NOE values (0.85-0.86), indicative of a well-ordered ∼22kDa domain. MD2 exhibited a slightly longer apparent correlation time (13.8 ± 1.16 ns), despite a similar domain molecular weight, suggesting a slightly increased effective hydrodynamic radius. The relaxation analysis for MD1 demonstrated limited fast internal motions at ps-ns time scale, mostly attributed to the loop regions; for MD2 an overall modest reduction in heteronuclear NOE (0.82-0.84) that could signify the presence of slightly more internal dynamics in the latter. Although diffusion anisotropy was negligible for both domains, MD2 exhibited higher R₂ variance and a lower backbone assignment completion rate than MD1 (66% vs 91% for MD1), particularly within ligand-binding pocket, consistent with more μs-ms conformational exchange than MD1. Thus, while both structures are globally very similar, MD2 seems intrinsically slightly more dynamic than MD1 (Fig. 1c).

### MD1 and MD2 bind ADPr with similar affinity but different thermodynamic profile

To gain further insights into the binding affinity and the thermodynamic parameters of the MD1 and MD2 interaction with ADPr we performed isothermal titration calorimetry (ITC). Both MD1 and MD2 bind ADPr with an affinity of 5.4 ± 0.4 μM for MD1, and a somewhat weaker affinity of 8.4 ± 0.6 μM for MD2 (Fig. 2a). These dissociation constants for ADPr are quite similar to that observed for other viral and human macro domains at similar buffer conditions^17^.

**Figure 2.**
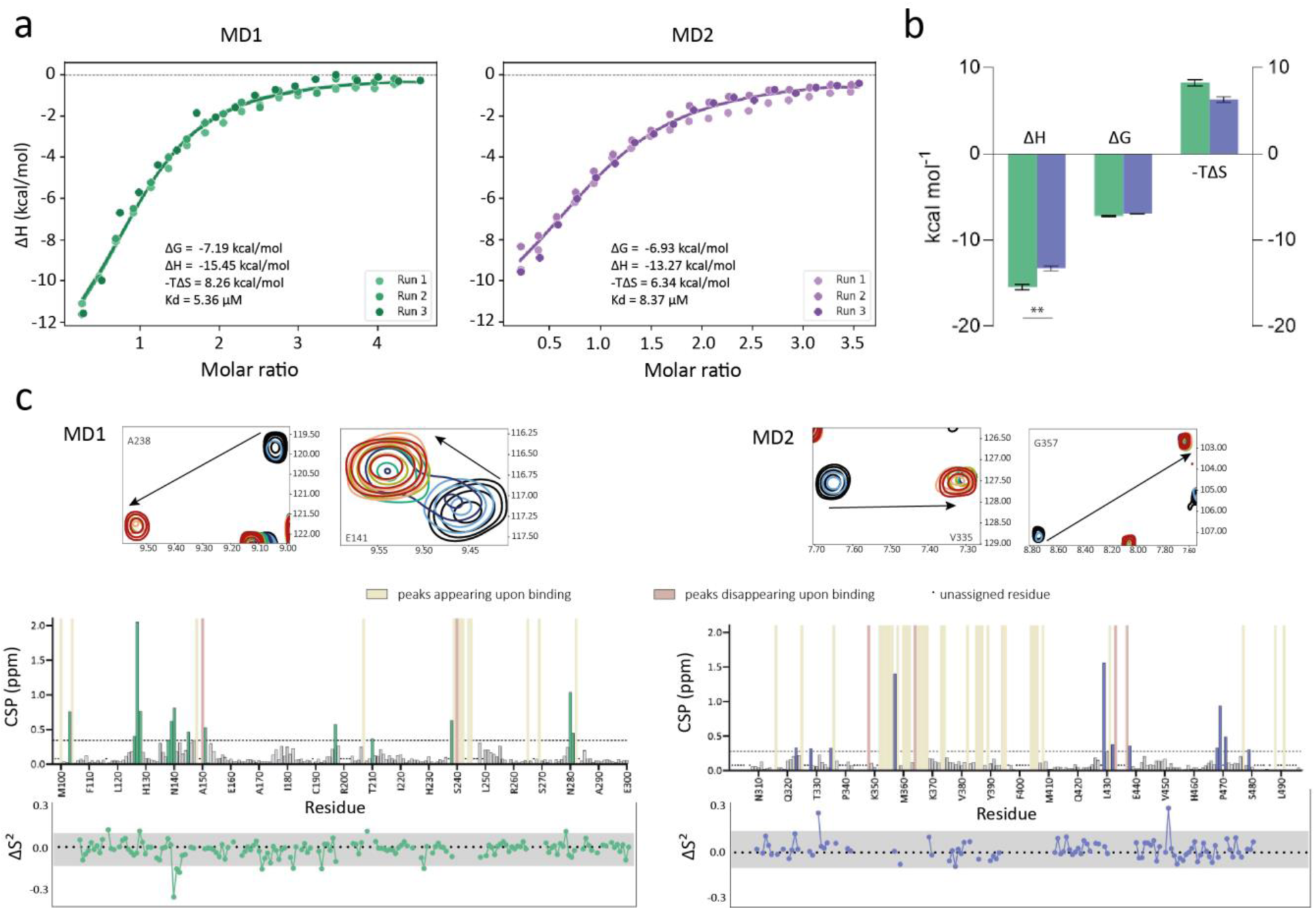
Biophysical and solution NMR characterization of PARP9 macro domain interactions with ADP-ribose. **a**. ITC analysis of PARP9-MD1 (green) and PARP9-MD2 (purple) binding to free ADPr. Three titrations per domain overlaid and fit to a single-site model. **b.** Comparative thermodynamic profiles (ΔH, ΔG, -TΔS) for MD1 (green) and MD2 (purple). Error bars denote 95% confidence interval; ** p<0.01. **c.** Spectral changes and residue-specific NMR Chemical Shift Perturbations (CSPs) for ¹⁵N-labeled PARP9-MD1 (data adapted from Moschidi, Fourkiotis et al.^18^) and MD2 upon titration with ADPr to saturation (1:5 molar ratio). Dashed lines represent significance thresholds (mean + 1 SD). Beige or pink bars indicate resonances that appear or disappear upon binding, respectively; black dots mark unassigned residues. Insets show representative expansions of the ¹H-¹⁵N HSQC spectra tracking the specific trajectories of selected residues (e.g., A238, E141, V335, and G357) in the free (black) and bound (blue/green/red) states. The bottom panel shows changes in backbone amide order parameters (ΔS² = S²bound - S²apo) versus residue number. Grey bands indicate the significance threshold (mean value with two standard deviations). Positive values indicate rigidification, negative values – decreased spatial restriction in presence of ADPr.

Although the K_D_ values, and henceforth the free energy of binding, ΔG, of MD1 and MD2 are similar, their thermodynamic profiles differ, under our assay conditions. The active MD1 binding is enthalpy driven (ΔH = −15.45 ± 0.31 kcal/mol) offset by a sizeable entropic penalty (-TΔS = 8.26 ± 0.34 kcal/mol), whereas the inert MD2 binds with significantly weaker enthalpy (ΔH = −13.27 ± 0.29 kcal/mol) and a smaller entropic cost (-TΔS = 6.34 ± 0.33 kcal/mol), resulting in similar Gibbs free energies of binding, (ΔG) of −7.19 ± 0.04 kcal/mol for MD1 and −6.93 ± 0.04 kcal/mol for MD2. The lower entropy contribution in MD2-ADPr complex may suggest that the ADPr ligand exhibits higher conformational freedom in the MD2 binding cavity than in the MD1 pocket (Fig.2b).

### ADPr binding causes limited structural changes in solution

To better understand ADPr binding to the two MDs, we performed NMR titration experiments to map spectral changes occurring upon binding. The binding to ADPr appeared to be in the slow exchange NMR time-scale for both domains (Fig. 2c), as indicated by decreasing backbone amide signal intensities with concomitant signal increases for the same amino acid at different location. In MD2, ADPr binding enabled assignment of numerous residues (31 residues, ∼16% of the whole sequence) that appeared broadened beyond detection in the apo state, suggesting that, particularly in known adenosine- and phosphate-coordinating loops, internal μs-ms dynamics is discarded upon ligand binding. The same but less pronounced effect was observed for MD1, where twelve additional residues were assigned in the bound state, with most of them residing in the loop between β6-α4 (residues 240-245), which is known as the phosphate binding loop (Fig. 2c).^18^

The ps-ns backbone dynamics were quantified from ^15^N R₁, R₂, and {^1^H}-^15^N NOE data. ADPr binding caused minimal changes in global tumbling for either domain (Δτc ≤ 0.4 ns), indicating that ligand engagement does not affect overall domain compactness. In MD1, the S² profile remained largely unchanged upon binding (Fig. 2c bottom), except for residue Asn140 and its neighboring residues Glu141 and Asp142, all localized within the catalytic loop. Instead of the expected rigidification that is frequently seen upon ligand binding, the observed slight local reductions in S² near the catalytic loop likely reflect subtle conformational adjustments required for ADPr recognition, whereas the adenosine-binding β2-β3 region remains rigid, consistent with a pre-organized binding site. Notably, residues within the phosphate-binding loop could be assigned after ADPr engagement, indicating ligand-induced rigidification. These conserved structural motifs together form the canonical NAD⁺-derivative binding pocket observed in both viral and human macro domains^3,19–21^. In summary, MD1 maintains a highly rigid backbone in both states, with ADPr binding inducing only minor, localized motions around the catalytic and phosphate-binding loops (Fig. 2c).

MD2 exhibits slightly increased the average S² values in the bound state compared to the apo state, indicating induced overall rigidification. Additionally, residues distal from the binding pocket (A331 and L451) seem to show increased S² values, which may suggest that a binding-induced decrease in flexibility propagates beyond the immediate interaction site and may reflect coordinated “breathing” motions that facilitate substrate recognition (Fig. 2c). Collectively, these data show that ADPr binding induces only subtle, site-specific adjustments in both MD1 and MD2, which seem unlikely to fully explain their different roles as ‘eraser’ vs ‘reader’ in PARP9.

### MD2 binds ADPr in two anomeric forms and shows greater binding-site plasticity than MD1

To detect potential structural differences that underline the different activities of MD1 and MD2, we determined X-ray crystal structures of both macro domains in complex with ADPr at resolutions of 1.44 Å and 1.30 Å, respectively (Fig. 3). At first glance, no major structural rearrangements are observed upon binding. The overall shape and binding site of both proteins remain similar (Fig. 3a). However, the accommodation of the ligand did induce larger displacement in the binding pocket of MD2 compared to MD1, as shown by RMSD values between the apo and ADPr-bound states, mapped in the worm thickness in Figure 3b. The binding site residues also correspond to high chemical shift perturbations (CSPs) observed upon NMR binding (Fig. 3c)

**Figure 3.**
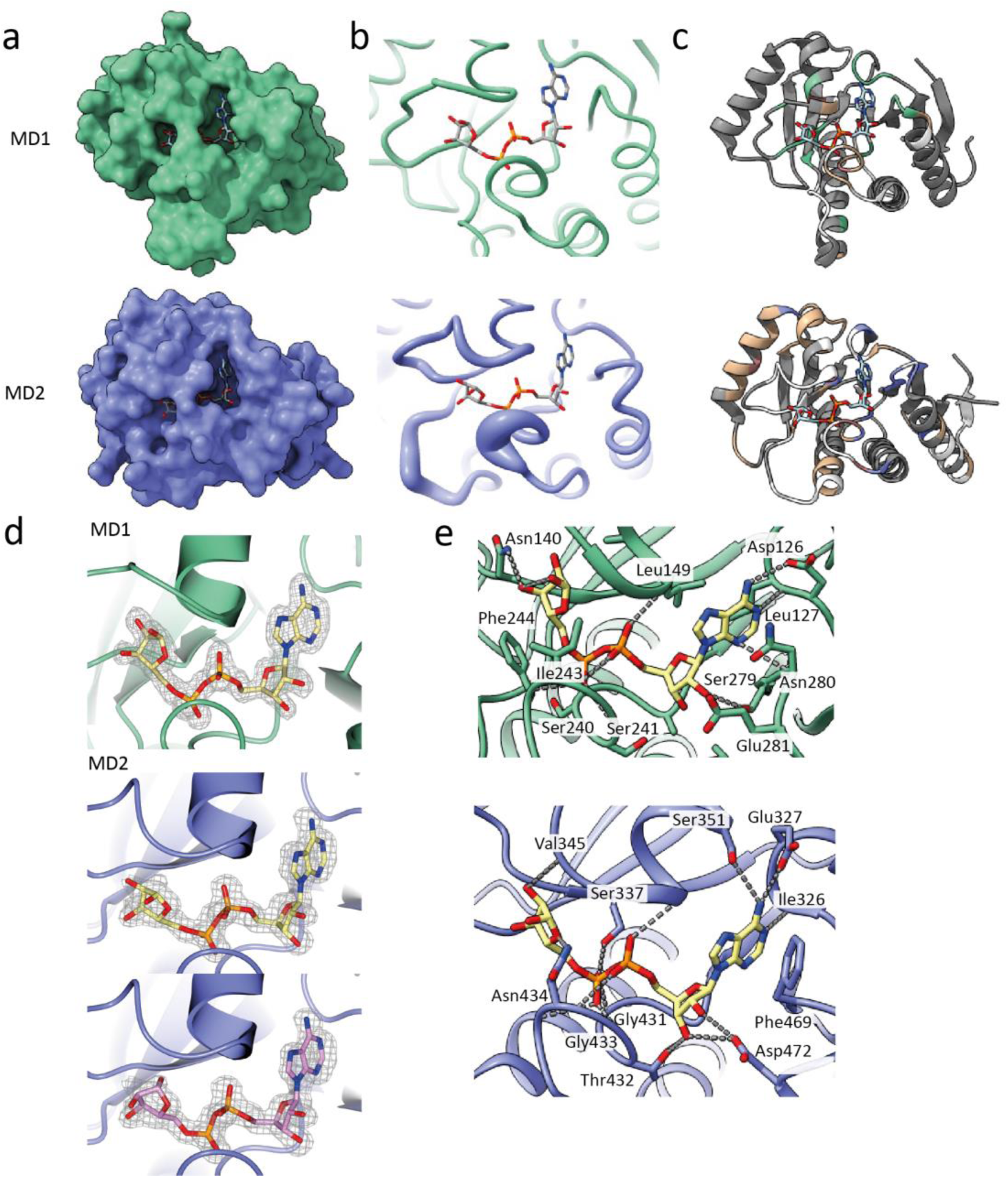
Crystal structures, conformational changes, and active-sites coordinating networks of PARP9 macro domains in complex with ADP-ribose. **a.** Solvent-accessible surfaces of ADPr-bound MD1 in green (9QYE) and MD2 in purple (9QYF) with ligand shown in sticks. **b.** Local backbone RMSD values between the apo and ADPr-bound structures mapped onto the cartoon models as worm thickness, showing greater active-site reorganization in MD2. **c.** NMR chemical shift perturbations (CSPs) from Figure 2c mapped onto the ADPr bound structures (white – unassigned residues; beige – residues assigned only in bound state; pink – residues assigned only in unbound state; green/purple – significant CSPs for MD1/MD2). **d.** 2Fo-Fc electron density maps (contoured at 1.0σ) of ADPr in MD1 and MD2 structures. In MD1, ADPr adopts a single well-defined conformation, in MD2 two anomeric states for the distal ribose were modelled. **e.** Close-up of ligand-coordinating networks in MD1 (top) and MD2 (bottom). Hydrogen bonds are denoted by dashed lines.

The ligand binding conformations are extremely well supported by the electron density (Fig. 3d), and interestingly, the ADPr molecule exists in two anomeric forms in MD2, but not MD1 (Fig. 3d, Supplementary Fig. 3). The binding mode is broadly conserved among MDs, with the α-phosphate, the distal ribose, and the adenosine being held in place by several residues. However, in the active MD1 structure Asn140 in the catalytic loop is binding the O3 of the ribose close to the scissile bond (Fig. 3e), which is not seen for its homologous amino acid N339 in the MD2.

### Family-wide comparison pinpoints specificity determinants for mutant design

Comparing our structures to the broader family of macro domains (Fig. 4a), offers additional insight. While adenosine, proximal ribose and phosphate moieties can be stabilized by various residues and display slight positional shifts across different proteins, for example, the adenosine position differs up to 4.9 Å between PARP9 MD1 and PARP14 MD1. The distal ribose consistently adopts the same position in all active proteins (such as PARP9 MD1) but is displaced in the inactive proteins (such as PARP9 MD2) (Fig. 4b-c). This difference could stem, at least partly, from differences in pocket architecture: MD1, as well as other active hydrolase macro domains, contain a conserved aromatic phenylalanine residue (Phe244) in the loop between the secondary elements β6-α4, which likely exerts steric and π-electron repulsion to the ADPr ligand. A previous study reported that substituting this phenylalanine with alanine reduced the de-MARylation activity of PARP9 MD1 by approximately 50%^14^. The corresponding residue in MD2 is methionine (Met435), lacking the capacity to exert similar constraints. Based on these observations we proceeded to design two groups of mutants.

**Figure 4.**
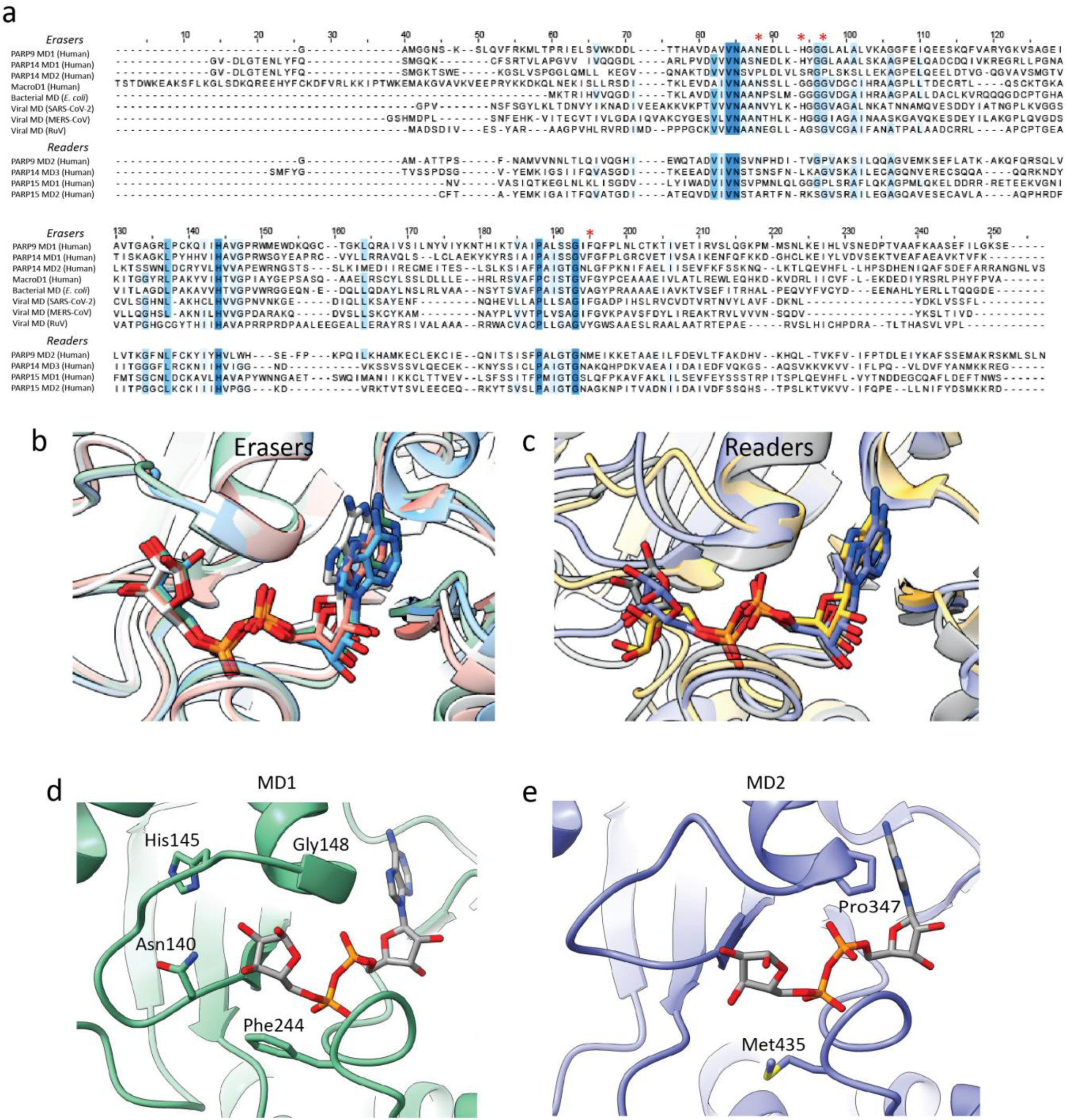
Sequence and structural comparison of PARP9 macro domains within the wider macro domain family and rationale for active-site variant design. **a.** Multiple sequence alignment of representative human, bacterial, and viral macro domains, grouped into “Erasers” (hydrolases) and non-catalytic “Readers”. Conserved motifs and critical residues are highlighted. **b, c.** Structural overlays of **b.** eraser and **c.** reader macro domains bound to ADPr. Erasers: PARP9-MD1 (9QYE) in green, bacterial MD (5FUD) in white, viral MD (6YWL) in blue, human MacroD2 (4IQY) in pink^22–24^. Readers: PARP9-MD2 (9QYF) in purple, PARP14-MD3 (4ABK) in yellow, macroH2A1.1 (3IID) in grey ^25,26^. **d, e.** Active-site close-ups of **d.** MD1 (green) and **e.** MD2 (purple), defining the structural blueprint for rational variant design. Mutation targets in MD1: Phe244 (positioning the distal ribose), Gly148 (loop flexibility), His145 and Asn140 (catalysis). Mutation targets in MD2: Pro347 and Met435.

In the first group of mutants, we attempted to abolish the MD1 catalytic function. We targeted the catalytic relevant Asn140, described before for its function^27^, and introduced mutations to alanine (N140A), to aspartate (N140D) to probe the importance of the charge, and to serine (N140S). The latter was chosen based on the TARG1 macro domain which has a structurally similar active site but is smaller by one β-strand and one α-helix^28^. Additionally, as indicated above, we designed a F244I mutant to potentially disrupt the positioning of the distal ribose that allows hydrolysis. We also mutated H145, which was shown to be important for water coordination in PARP14 MD1 and coronaviral MDs, to arginine and tyrosine (H145R, H145Y). Here, arginine is a mutation that probes the hydrogen bond making ability of histidine, while tyrosine could potentially substitute for its π -staking properties. Finally, a G148P was designed to probe the role of catalytic loop flexibility, as the introduction of a proline residue at this position is expected to impose conformational constraints on the loop (Fig. 4d).

The second group of mutants aim to introduce mutations in MD2 that appear to stabilize the distal group in active proteins (Fig. 4b) and thus engineer catalytic activity. These substitutions were designed to mimic the interaction pattern observed in MD1 and other “eraser” homologs (Fig. 4a-b). We introduced a P347G mutation to mimic G148 in MD1, reintroducing the flexibility characteristic of the glycine-rich loop found in MD1 and other active macro domains. We also introduced a M435Y mutation, as all active macro domains carry aromatic residue at this position, seemingly pushing the distal ribose upwards and acting as a “lid”, leaving no space for it to deviate from the correct position (Fig. 4e).

### Single-point mutations disrupt MD1 catalysis and unlock latent activity in MD2

All tested substitutions at positions 145 and 148 and the N140A/N140S mutants of MD1 resulted in unstable protein and could not be purified. The N140D and F244I mutants of MD1 could be expressed and purified, similar to both MD2 variants, P347G and M435Y. We then tested these four mutants for different functional aspects.

The N140D mutant of MD1 showed ∼4-fold increased affinity for ADPr (K_D_ = 1.44 ± 0.2 μM) (Fig. 5a) compared to wild-type, but its protein de-MARylation activity was reduced. Notably, a homologous Asn-to-Asp substitution in the SARS-CoV-2 macro domain (N40D) was previously reported to abolish catalytic activity entirely, suggesting significant differences in their active-site microenvironment could perturb catalysis (PARP9) or fully disable it (SARS-CoV-2)^27^. In contrast to the protein substrate, however, RNA de-MARylation activity of the N140D MD1 was completely lost. The F244I mutant displayed a markedly reduced affinity in the ITC experiment (Fig. 5a) and considerably impacted the de-MARylation activity of MD1 towards protein and RNA substrates (Fig. 5b).

**Figure 5.**
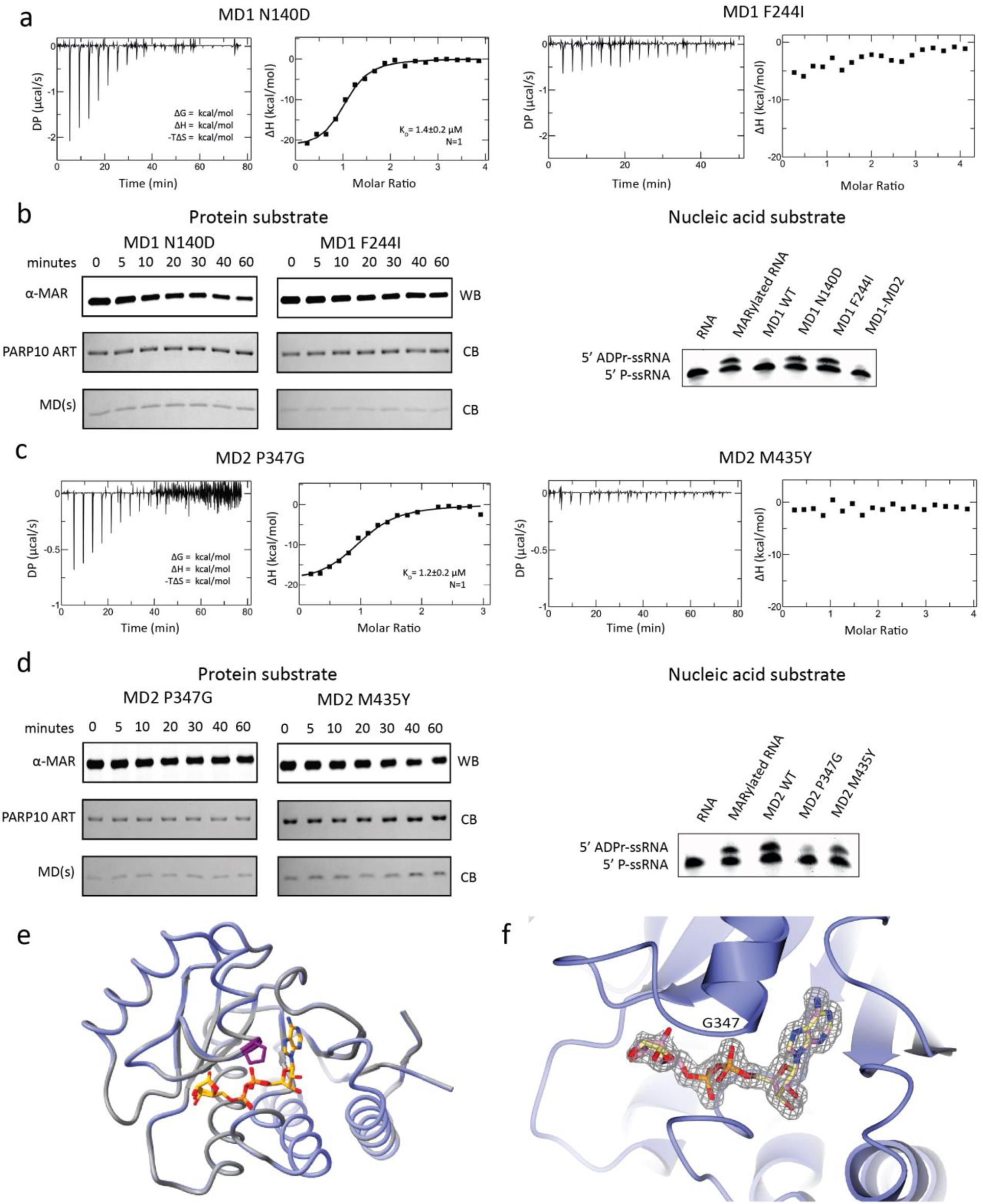
Functional validation, thermodynamic profiling, and structural characterization of PARP9-MD1 and MD2 active-site variants. **a.** ITC thermograms (left) and integrated binding isotherms (right) for MD1 mutants N140D and F244I with ADPr. Extracted dissociation constants (KD) and single-site model fits are displayed. All titrations were performed in triplicates. **b.** MD1 variants activity. De-MARylation of MARylated PARP10 ART monitored via Western blot (left, time-dependency over 60 minutes) and urea-PAGE gel-shift assay evaluating activity toward the 5’ ADPr-ssRNA substrate (right, 30 minutes endpoint). MD1 WT and MD1-MD2 WT are active against ssRNA substrate; both mutants are not. **c.** ITC thermodynamic profiling of MD2 variants P347G and M435Y with ADPr, providing corresponding thermograms, binding curves, and calculated KD values. All titrations were performed in triplicates. **d.** MD2 variants activity by Western blot (left) and urea-PAGE gel-shift assay (right), showing an RNA-specific gain of function for P347G. **e.** Overlay of MD2 WT bound to ADPr in grey (9QYF) and MD2 P347G bound to ADPr in purple (32HC). Residues 347 are shown in dark magenta. ADPr molecules of the mutant structure are shown in orange sticks. **f.** Close up of ADPr inside P347G binding pocket. The 2Fo-Fc electron density map (contoured at 1.0σ) surrounding the ligand is shown as a grey mesh.

The P347G mutant of MD2 showed a 7-fold tighter binding (K_D_ = 1.17 ± 0.2 μM) for ADPr, compared to wild-type, while the M435Y mutation drastically reduced affinity (Fig. 5c). Neither MD2 mutant displayed detectable de-MARylation activity toward MARylated PARP10 ART (Fig. 5d). Strikingly, however, the P347G variant exhibited detectable activity toward RNA de-MARylation (Fig. 5d, Supplementary Fig. 4), representing a clear gain of function for a domain previously considered catalytically inert. Notably, the P347G/M435Y double mutant was completely inactive in this assay (Supplementary Fig. 4), demonstrating that M435 and P347 do not independently contribute to catalytic competence. Potentially, the gain of function conferred by P347G in MD2 depends on a precise configuration of the catalytic loop, which could be disrupted by the additional M435Y substitution.

We previously pointed out that MD2 binds with weaker enthalpy and a smaller entropic cost compared to MD1, even though the ADPr binding affinities are similar. To find out if this could be related to eraser vs reader functionality, we characterized the binding thermodynamics of the MD2 P347G mutant. Indeed, ADPr binding to P347G shows higher enthalpy (ΔH = −19.8 ± 0.7 kcal/mol) than MD1 or MD2 wild type (ΔH of −15.45 ± 0.3 and −13.27 ± 0.3 kcal/mol, respectively) and pays a higher entropic penalty (-TΔS = 11.7 kcal/mol) than MD1 or MD2 wild type (-TΔS of 8.26 and 6.34 kcal/mol respectively) making it more MD1-like in the context of its thermodynamic profile. We therefore conclude that the reason this single point mutation is sufficient to unlock latent catalytic potential in MD2 is owed to its conformational dynamics. Moreover, we suggest that the selective activity toward MARylated RNA, but not MARylated protein, could be because it preferentially induces a conformational plasticity compatible to accommodate and hydrolyze ADPr units on nucleic acids.

To further test this hypothesis, we determined X-ray structures of MD2 P347G mutant both in apo and ADPr-bound states. Comparison with the wild type MD2 revealed no conformational changes, indicating that the mutation neither imposes local or large protein conformational changes, nor alters the shape of the binding site (Fig. 5e-f, Supplementary Fig. 6a). B-factor analysis showed slightly increased flexibility of the mutated region, consistent with our previous hypothesis. The mutant binds ADPr in a manner resembling that observed for the wild type. Analogous to the wild type, two conformations of ADPr could be modelled, although this was based on our prior knowledge, as the electron density could support either conformation with only small positive difference density for the respective second conformation. The distal ribose appeared to be even more flexible in the mutant structure, which is noticeable by the poorer electron density and somewhat elevated normalized B-factors (Supplementary Fig. 6b). The newly introduced residue establishes an extra interaction with the phosphate group of ADPr, which could account for the observed change in enthalpy of binding (±6.5 kcal/mol).

The data presented herein highlighted two distinct differences between MD1 and MD2, namely the glycine-rich loop and the lower region of the binding pocket which includes Phe244 in MD1. The mutant structures provide further insight into the respective roles of these elements. These data suggest that the glycine-rich loop itself is not the main determinant of the appropriate orientation of the distal ribose. Instead, it seems to provide sufficient adaptability for the loop to accommodate the ribose in the catalytically correct orientation. This may also help to explain the observed differences between mutants’ activity for MARylated protein and RNA substrate. ADP-ribosylated RNA might promote the proper ribose conformation, while the glycine permits its accommodation.

## Concluding remarks

The two macro domains of PARP9 present a striking paradox: they share the canonical α/β/α fold, near-identical binding pocket shapes, and comparable picosecond-nanosecond dynamics, yet one is an active eraser and the other a catalytically silent reader. We find that this divergence is not encoded in the catalytic machinery, as the catalytic asparagine (Asn140/Asn339) is structurally conserved in both macro domains and, more broadly, across many other macro domains, but rather in the details of the architecture and adaptability of the environment that positions the distal ribose of ADPr. MD1 is pre-organized, paying an entropic cost to hold the distal ribose in a catalytically productive geometry, whereas MD2 remains flexible and binds ADPr without committing it to a reactive pose. The distinct thermodynamic signatures of free-ADPr binding, despite similar affinities, capture this difference directly.

The broader principle is that the two states are separated by so little: a single proline-to-glycine substitution in the catalytic loop (P347G) is enough to confer de-MARylation activity on the otherwise inert MD2, indicating that the reader-eraser distinction across the macro domain family is a subtle, tunable and evolutionarily accessible property rather than a fixed structural divide.

Two essential questions remain unanswered from our studies, namely why the restored activity in MD2 is selective for ADP-ribosylated RNA over protein, and how the eraser-reader division operates within full-length PARP9 in cellular processes. Resolving these will clarify how PARP9 couples ADP-ribose recognition and removal to its roles in RNA-virus sensing, interferon signaling, and the DNA-damage response, and whether its two distinct ADPr binding pockets can be exploited as selective therapeutic targets in malignancies where PARP9 is implicated.

## Materials and Methods

### Construct Design

Residues 102-300 and 305-497 of full-length (fl) human PARP9 (UniProt entry: Q8IXQ6) were used to generate constructs for hPARP9 MD1 and MD2, respectively. A double-domain construct containing both MDs comprised residues 102-497 of fl-hPARP9. A synthetic, codon-optimized gene of fl-hPARP9 (GenScript) was amplified by PCR, and the sequences encoding MD1, MD2, and MD1-MD2 were cloned into a pETM-41 vector. The PARP10 construct (residues 800-1025) was obtained from GenScript already cloned into a pGEX-4-T1 plasmid.

For wild-type hPARP9 MDs, the primer sequences were as follows: MD1 forward 5’-CATGCCATGGGAGGTAACAGCAAGAGCCTG-3’ and reverse 5’-ATAGTTTAGCGGCCGCTTAGCTACGTTTCGC-3’; MD2 forward 5’-CATGCCATGGAGACCACCCCGAGCTTC-3’ and reverse 5’-ATAGTTTAGCGGCCGCTTACACGCTATAGTTGTTC-3’. For the MD1-MD2 construct, the forward primer of MD1 and reverse primer of MD2 were used. Primers for mutant constructs are listed in Supplementary Table 1. All resulting constructs were verified by DNA sequencing. The expressed proteins carried an N-terminal His₆-MBP tag followed by a TEV protease cleavage site, leaving four artificial N-terminal residues (GAM[G/A]) in the final studied proteins.

### Protein expression and uniform ^15^N and ^15^N/^13^C labeling

Plasmid encoding the PARP9 MD1-MD2 was transformed into *Rosetta™ 2(DE3) pLysS* cells. An LB pre-culture was grown at 37 °C with shaking (180 rpm) for 14-16 h, and subsequently used to inoculate 0.5 L of M9 minimal medium (48 mM Na₂HPO₄, 22 mM KH₂PO₄, 8 mM NaCl) supplemented with 0.5 g ¹⁵N-labeled NH₄Cl, 2 g unlabeled or ¹³C D-glucose, 1 mL of a stock solution containing 0.5 mg/mL biotin and 0.5 mg/mL thiamine, 0.5 mL of 1 M MgSO₄, 0.15 mL of 1 M CaCl₂, and 1 mL of trace elements (40 mM HCl, 50 mg/L FeCl₂·4H₂O, 184 mg/L CaCl₂·2H₂O, 64 mg/L H₃BO₃, 18 mg/L CoCl₂·6H₂O, 4 mg/L CuCl₂·2H₂O, 340 mg/L ZnCl₂, 710 mg/L Na₂MoO₄·2H₂O, 40 mg/L MnCl₂·4H₂O). For antibiotic selection, 50 µg/mL kanamycin and 34 µg/mL chloramphenicol were used. Cultures were incubated at 37 °C, 180 rpm until OD₆₀₀ reached 0.6-0.8, at which point expression was induced with 1 mM IPTG at 25 °C.

### Protein purification

Harvested cells expressing His_6_-MBP tagged PARP9 MD1-MD2 were lysed by sonication in 50 mM Tris pH 7.0, 1 M NaCl, 10% glycerol, 10 mM imidazole, 2 mM DTT, DNase I, and protease inhibitors. Cleared lysate was loaded onto a HisTrap HP column (Cytiva) and eluted across an imidazole gradient (40 to 400 mM). The eluate was buffer exchanged into 50 mM Tris pH 8.0, 300 mM NaCl, 2 mM DTT using a 10 kDa MWCO filter (Amicon), treated with TEV protease overnight at 4 °C, and passed through a secondary IMAC column to capture the cleaved tag. Cleaved PARP9 MD1-MD2 was finalized by size exclusion chromatography on a Superdex 200 10/300 GL column (Cytiva) equilibrated in 10 mM HEPES pH 7.0, 20 mM NaCl, 2 mM DTT, 2 mM EDTA. Pure fractions were pooled, concentrated, and stored at −80 °C.

All the different mutants followed the purification protocols of the corresponding wt. The expression and purification of hPARP10 ART domain is mentioned elsewhere^17^ and for the single domain constructs of PARP9, MD1 and MD2 will be described in another publication^18^.

### Isothermal Titration Calorimetry - ITC

ITC experiments were performed at 25 °C using a MicroCal PEAQ ITC instrument (Malvern, UK). All proteins and ligands were prepared in 50 mM HEPES at pH 7.0, 50 mM NaCl, and 2 mM EDTA. ADPr was titrated into 17 to 18.3 µM of PARP9 MD1 and 22 to 22.4 µM of PARP9 MD2 using a stock solution of 0.4 mM (0.8 mM for the MD1-MD2 double domain construct). Each titration consisted of 19 injections and was performed in triplicate. For unstable mutants, in high concentrations, that could not be retained in the cell during titration, reverse titrations were performed where ADPr was loaded into the cell and the macro domain proteins were loaded into the syringe at a concentration of 100 µM.

Initial isothermal titration calorimetry thermograms were done using MicroCal PEAQ ITC Analysis Software version 1.41. Binding curves for single domain constructs were fitted using a standard one site binding model, and for reverse titrations, the ligand in cell configuration was utilized. The stoichiometry N was fixed to 1 for all experiments, reflecting the known 1 to 1 interaction between the macro domain and ADPr. To account for the enthalpy of titrant dilution, corresponding control experiments where ligand was titrated into buffer, or buffer into protein for reverse titrations, were subtracted from the experimental data prior to curve fitting.

For MD1 ADPr and MD2 ADPr, a single site Wiseman binding model with N fixed at 1 was fitted globally across all three replicates per protein, with ΔH and K_D_ shared as common parameters. Normalized heats were recomputed from raw ΔQ. Uncertainties in ΔG = RTln(K_D_) and -TΔS = ΔG - ΔH were propagated from the covariance matrix of the nonlinear least squares fit. All confidence intervals are reported at the 95% level using a t distribution with 52 degrees of freedom per protein. This was corroborated by two model free analyses that made no assumption about the shape of the binding isotherm: pointwise Welch t-tests with Benjamini Hochberg correction on interpolated heat vectors identified significant differences at low molar ratios, and a Welch t-test on PC1 scores from principal component analysis of the raw heat curves, which captured 84% of total variance, showed complete separation between MD1 and MD2 with a p value of 0.0085. The code for analysis was implemented by Claude Opus 4.8 and is submitted to Zenodo (DOI: 10.5281/zenodo.21337531).

### Protein MARylation/De-MARylation assays

MARylation assays were performed as previously described^17^ using as a substrate the GST-hPARP10 ART domain. Briefly for the MARylation reaction the hPARP10 ART domain was incubated with β-NAD (β-NAD^+^, N1636 Sigma Aldrich), 1:100 molar ratio, for 20 min at 37 °C in 50 mM HEPES pH 7.5, 150 mM NaCl, 0.2 mM DTT, 0.02% IGEPAL® (CA-630). The MARylated substrate was then purified by size-exclusion chromatography (SEC) using a Superdex™ 200 Increase 10/300 GL column (Cytiva).

De-MARylation assays were performed as a single continuous reaction at 30 °C and 250 rpm. The mixture contained 1 μM MD and 1 μM MARylated substrate in a buffer of 50 mM HEPES pH 7.5, 150 mM NaCl, 0.2 mM DTT, 0.02% IGEPAL® CA-630, and 2% DMSO, supplemented with or without specified ratios of ADPr. To monitor the time course, individual fractions were sampled at the indicated time points and quenched immediately by the addition of 4× LDS buffer. The samples were heated at 70 °C for 3 min, centrifuged at 14,000 rpm and 4 °C for 5 min, and then separated on a 12% Bis-Tris polyacrylamide gel. Each experiment was performed three times.

### RNA ADP-ribosylation/de-MARylation

For RNA MARylation, 10 μM single-stranded RNA (5′P-GUUUCGGAUCGACGC-3′)^13^, obtained from Microsynth, was incubated with 10 μM GST-tagged PARP10 ART domain and 1 mM β-NAD⁺ (β-NAD^+^, N1636, Sigma-Aldrich) for 90 minutes at 37 °C in reaction buffer containing 50 mM HEPES pH 7.5, 50 mM NaCl, 1 mM DTT, 5 mM MgCl_2_. Furthermore, PARP10 ART was digested with 20 U Proteinase K (New England Biolabs) at 37 °C for 30 min. After that, the modified RNA was also further purified using an RNA purification kit (Monarch, NEB #2030). The purified 5’ADP-ribosylated RNA was used in the indicated assays.

Hydrolase assays were performed by incubating 10 μM ADP-ribosylated RNA with 1 μM of the respective macro domain (MD) protein in the ADP-ribosylation buffer, for 30 min. While the time-course assays were performed, in triplicates, by incubating 1 μΜ ADP-ribosylated RNA with 0.18 μΜ of each MD and samples were taken at the indicated time points. Reactions for hydrolase assays were terminated by the addition of RNA loading dye containing 80% (w/v) formamide, 10 mM EDTA (pH 8.0), 1 mg/mL xylene cyanol FF, and 1 mg/mL bromophenol blue. Samples were heated at 95 °C for 3 minutes before loading onto pre-run denaturing urea-polyacrylamide gels (8 M urea, 20% acrylamide:bisacrylamide [19:1], 0.25% APS, 0.5% TEMED).

Electrophoresis was performed in 1× TBE buffer (89 mM Tris, 89 mM boric acid, 2 mM EDTA, pH 8.3) at constant 140 V and 4 °C. Oligonucleotides were visualized using the ChemiDoc Imaging System (Bio-Rad) following staining with SYBR™ Gold nucleic acid gel stain (Invitrogen).

Band intensities were quantified by densitometry using uniform exposure settings within each gel. Background subtraction was applied uniformly across lanes. The fraction of ADP-ribosylated RNA remaining at each time point was calculated relative to the total signal in the corresponding lane and normalized to the signal at time zero for each reaction series.

The initial rate of substrate decay (k) was determined as previously described for protein MARylated substrates^17^.

### Immunoblotting

Proteins were transferred onto a polyvinylidene fluoride (PVDF) membrane (0.45 μm pore size; Immobilon-P, Merck Millipore) using a semi-dry transfer system (Bio-Rad). Membranes were blocked for 1 hour at room temperature in Tris-buffered saline (TBS, pH 7.5) containing 0.1% Tween-20 and 5% (w/v) non-fat dry milk. Blots were then incubated with anti-mono-ADP-ribose reagent (MABE1076, Merck Millipore) overnight (14-16 hours) at 4 °C.

Detection was performed using an HRP-conjugated anti-rabbit IgG secondary antibody (7074, Cell Signaling Technology). Protein bands were visualized using a ChemiDoc Imaging System (Bio-Rad), and band intensities were quantified using Image Lab software (Bio-Rad).

The initial rate of substrate decay (k) was determined as previously described.^17^

### Crystallization, data collection, data processing and refinement

Conditions for crystal growth of MD1 and MD2 were screened using commercial screening plates BCS, JCSG, PACT and Morpheus (Molecular Dimensions) with the vapor diffusion sitting drop method. Protein stock solutions were used in concentrations of 10 mg/mL in free form and bound to ADPr. The drops were dispensed by a Mosquito Xtal3 (SPT Labtech) with a volume of 200 nL and 50 : 50 protein-to-screening condition ratio. Subsequently, the conditions of the initial hits were further optimized, varying component concentrations. For MD1 in free form an additive screen was used (Hampton Research). The final conditions yielding suitable crystals were: for free MD1 2.4 M Sodium malonate pH 7, 4% PPG P400; for MD1 bound to ADPr 0.1 M Imidazole/MES buffer pH 6.5, 20 % v/v PEG 500MME, 10 % w/v PEG 20000, 0.12 M Monosaccharides mix; for free MD2 0.1 M MIB pH 7, 25% w/v PEG 1500; for MD2 bound to ADPr: 0.1 M Succinic acid, 15 % PEG 3350; for free MD2 P347G 0.1 M NPS, 0.1 M Imidazole/MES buffer pH 7, 12.5% v/v MPD; 12.5% PEG 1000; 12.5% w/v PEG 3350: and for MD2 P347G bound to ADPr 0.2 M Sodium nitrate, BTP buffer pH 8.5, 20 % w/v PEG 3350.

Crystals were mounted on cryo-loops and vitrified in liquid nitrogen; 30% glycerol was used as a cryo-protectant. The X-ray diffraction data were collected at the European Synchrotron Radiation Facility (ESRF) with the wavelength of 0.97 Å. Data were processed with XDS^29^ or DIALS^30^ and scaled with AIMLESS^31^. The data for all MD2 structures were truncated based on CC_1/2_, completeness, and |I/σI|. For MD2 bound to ADPr phasing was done with the molecular replacement with MOLREP^32^ using structure with the PDB ID 5AIL as a model. Subsequently, MD2 bound to ADPr was used as a starting model for MD2 free, and both MD2 P347G structures. For MD1 free form the *in silico* model from AlphaFold with accession code AF-Q8IXQ6-F1 was used as a starting model and later our experimental model was used for phasing for MD1 bound to ADPr. The model building was carried out using Coot and refinement was done using REFMAC^33^ from CCP4i2 suite^34^. The ADPr was built using Coot into electron density inside the binding pockets of MD1 and MD2. All structures bound to ADPr were refined using anisotropic B-factors, the rest of the structures were refined with TLS refinement, additionally for MD2 WT and mutant in free form NCS restraints were applied during refinement. The PDB-REDO server was used for further refinement^35^. Final structures were assessed with MolProbity^36^ and Rama-Z analysis was performed using Tortoize web server^37^. All data collection and refinement statistics can be found in Table 1.

**Table 1.**
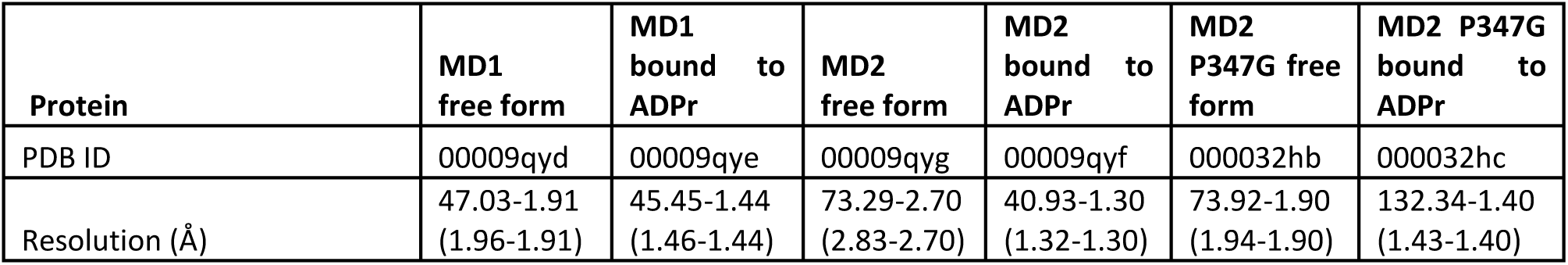
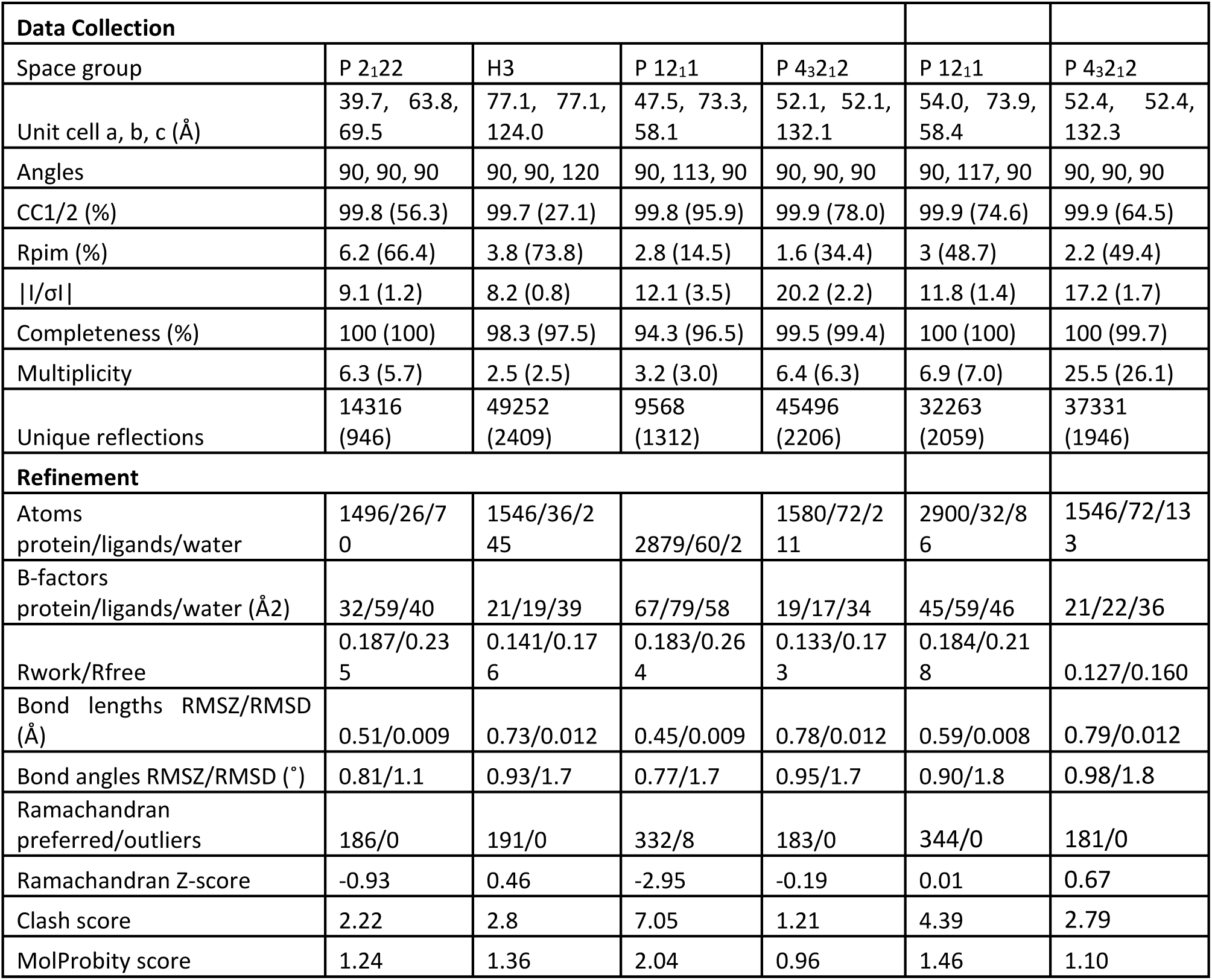
Data collection and refinement statistics.

### Chemical shift perturbation

For ADPr titration experiments, wild-type PARP9 ^15^N-MD1 was prepared in a buffer containing 10 mM HEPES (pH 7.1), 50 mM NaCl, 2 mM TCEP, and 2 mM EDTA. For ^15^N-MD2 the buffer consisted of 10 mM HEPES (pH 7.0), 20 mM NaCl, 2 mM DTT, and 2 mM EDTA. All NMR samples contained 10% D₂O and 0.25 mM DSS (4,4-dimethyl-4-silapentane-1-sulfonic acid) as an internal reference. Spectra were measured at 25 °C using a Bruker Avance III HD four-channel 700 MHz NMR spectrometer equipped with a cryogenically cooled 5 mm ^1^H/^13^C/^15^N/^2^H Z-gradient TCI probe.

NMR titrations were performed by incrementally increasing the [protein]:[ligand] ratio (e.g., 1:0.25, 1:0.5, 1:1, 1:3, 1:5, 1:10), monitored by a ¹H-¹⁵N HSQC spectrum after each addition. Titrations were completed at a final ratio of 1:10. Combined amide chemical shift perturbations (CSPs) were calculated using the equation:

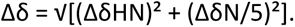

To identify significant CSPs, a threshold was defined by calculating the mean CSP and standard deviation (σ), iteratively excluding residues with values greater than 3σ at each step. The final threshold for meaningful CSPs was set as the mean plus one standard deviation^38,39^.

Earlier MD1 CSP data has been presented elsewhere^18^ and are shown alongside newly acquired MD2 CSP data (this study) for comparative analysis.

### 15N-relaxation studies

Sensitivity-enhanced TROSY-type pulse sequences were used to measure ^15^N backbone spin relaxation. A Bruker Avance III HD four-channel 700 MHz NMR spectrometer equipped with a cryogenically cooled 5 mm ^1^H/^13^C/^15^N/^2^H Z-gradient TCI probe was used to get three-dimensional pseudo-spectra.

Spectra were measured at 25 °C with a spectral width (SW) of 14 ppm over 2048 complex points in the ^1^H dimension and 44 ppm over 128 complex points in the ^15^N dimension. 16 transients (NS) were collected for measurements of longitudinal (*R*₁) and transverse (*R*₂) relaxation rates. Each relaxation experiment in the series was preceded by a relaxation delay (D₁) of 1.2 s. Ten relaxation delays (20, 60, 100, 200, 400, 600, 800, and 1200 ms), including repetitions at 60 and 600 ms, were used to calculate *R*_2_ relaxation rates. Ten Carr–Purcell– Meiboom–Gill (CPMG) delays of 16.96, 33.92, 67.84, 101.76, 135.68, 169.60, 203.52, and 237.44 ms were used to quantify *R*₂ relaxation rates, with repetitions at 33.92 and 203.52 ms.

TROSY-type experiments were used to obtain backbone {^1^H}-^15^N steady-state heteronuclear NOEs. Using NS = 16, two-dimensional spectra comprising NOE-enhanced and reference (unsaturated) datasets were obtained with D₁ = 3 s, NS = 16, SW(^1^H) = 14 ppm (2048 complex points), and SW(^15^N) = 44 ppm (128 complex points). The NOE was calculated as the ratio of peak intensities in saturated and unsaturated spectra.

TopSpin (Bruker BioSpin) was used to process all relaxation datasets, and Dynamics Center version 2.8.8 (Bruker BioSpin) was used for relaxation rate fitting, heteronuclear NOE calculation and analysis. Monte Carlo error propagation was used to determine the uncertainties in *R*₁ and R₂ values.

With a ^15^N chemical-shift anisotropy value (CSA) of −160 ppm and an NH bond length of 1.02 Å, generalized order parameters (*S*²) were extracted using Lipari-Szabo model-free analysis of the relaxation parameters in Dynamics Center^40,41^. Furthermore, 1000 random selections of the start parameters were used to improve model fitting. The average *R*₂/*R*₁ ratio of residues in well-structured regions was used to estimate the total rotational correlation time (τ_c_) and molecular tumbling anisotropy. Slower (μs-ms) conformational exchange was thought to be occurring in regions with abnormally high *R*₂ or low {^1^H}-^15^N NOE values (<0.65) and they were excluded from the calculation.

The crystal structures of each protein in their respective states were used to calculate the principal values of the rotational diffusion tensor, corresponding to the three eigenvalues Dxx, Dyy, and Dzz. The isotropic rotational diffusion coefficient (Diso) was then determined as one third of their sum:

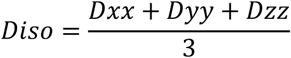

N-H bond vector orientations derived from the crystal structure of each domain/state were used in Dynamics Center (Bruker) to fit the rotational diffusion tensor against the experimental R₁/R₂ relaxation rates. The resulting anisotropies (Dzz/Dxx = 1.08 for MD1-ADPr, 0.97 for MD1-apo, 0.95 for MD2-ADPr, and 0.99 for MD2-apo) were close to unity in all four datasets, indicating essentially isotropic overall tumbling. A single global τ_c_ was therefore used per dataset in the subsequent Lipari-Szabo model-free analysis, rather than an axially symmetric or fully anisotropic diffusion model.

The τ_c_ was subsequently calculated using the relation:

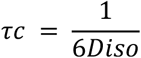

To discuss the influence of ADPr binding on macro domains we calculated ΔS2 calculated as: S²bound - S²apo. In each plot, the dotted horizontal line indicates the threshold applied to identify statistically meaningful dynamical changes, calculated based on the absolute values of Δ*S*^2^ using the mean (M) plus two times the standard deviation (SD):

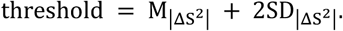

## Acknowledgements

We acknowledge EU-finding through the HORIZON-WIDERA-2022-TALENTS-01 ERA Chairs “ESPERANCE” project (GA 101087215 (DOI: 10.3030/101087215)), and the HORIZON-WIDERA-2023-ACCESS-04 Pathways to Synergies “MILESTONE” project (GA 101159708 (DOI: 10.3030/101159708)). We also acknowledge EU FP7 REGPOT CT-2011–285950—“SEE-DRUG” project for the purchase of UPAT’s 700 MHz NMR equipment. This work benefited from access to the NKI Protein Facility, an Instruct-ERIC center and was supported by iNEXT-Discovery project (No 871037, H2020, EC) and Instruct-ERIC (PID 37444). Work at the NKI has been supported by an institutional grant of the Dutch Cancer Society and of the Dutch Ministry of Health, Welfare and Sport. ESRF data collection was done under Dutch BAG proposals MX2526, MX2649 and MX2741. We thank Hans Wienk for critically reading the manuscript and Robbie Josten for crystallographic discussions.

## Author Contributions

N.K.F. performed most of the experiments, including crystallization, structure determination and design of mutants (jointly with A.C.), analyzed data, and wrote the initial draft of the manuscript. C.S.B. performed protein purification, functional assays, and data analysis, and edited the manuscript. A.C.T. contributed to conceptualization and data analysis and edited the manuscript. A.F. analyzed and validated thermodynamic data. K.P.K. carried out experimental work regarding the PARP9 MD2 expression and purification protocol and edited the manuscript. S.A.T. designed the initial MD1 and MD2 constructs and contributed to conceptualization. A.P. discussed experimental ideas and approaches, performed data analysis, and edited the manuscript. A.C. performed or supervised all crystallographic experiments and analysis, collated all data, substantially revised the manuscript, made all figures, and edited the final manuscript. G.A.S. conceived and supervised the study, provided resources and funding acquisition, acquired all NMR data, and edited the manuscript. All authors reviewed and approved the final manuscript.

## Accession numbers

PDB IDs: PARP9MD1 APO: 9QYD, PARP9MD1 ADPr: 9QYE, PARP9MD2 ADPr: 9QYF, PARP9MD2 APO: 9QYG, PARP9MD2 P347G APO: 32HB, PARP9MD2 P347G ADPr: 32HC

