## Supplementary figures for "So similar, yet so different: the paradigm of PARP9 macro domain paralogs"

**Supplementary Table 1.** Primers used for PARP9 macro domains mutants

| Mutant | Forward primer | Reverse primer |
|---|---|---|
| MD1 N140A | CGCGGCGGCGGAGGATCTGCTGCACGGTGG | TCCTCCGCCGCCGCGTTAACCACCGC |
| MD1 N140D | CGCGGCGGATGAGGATCTGCTGCACGGTGG | TCCTCATCCGCCGCGTTAACCACCGC |
| MD1 N140S | CGCGGCGAGCGAGGATCTGCTGCACGGTGG | TCCTCGCTCGCCGCGTTAACCACCGC |
| MD1 H145R | TCTGCTGCGCGGTGGCGGTCTGGCGCTGG | CCACCGCGCAGCAGATCCTCGTTCGCC |
| MD1 H145Y | TCTGCTGTATGGTGGCGGTCTGGCGCTGG | CCACCATACAGCAGATCCTCGTTCGCC |
| MD1 G148P | CGGTGGCCCGCTGGCGCTGGCGCTGGTG | GCCAGCGGGCCACCGTGCAGCAGATC |
| MD2 P347G | GATATTACCGTGGGCGGCGTTGCGAAAAGCATCC | GGATGCTTTTCGCAACCCCCCCCACGGTAATATC |
| MD2 M435Y | GCTGGGCACCGGTAACTATGAAATCAAGAAAGAG | CTCTTTCTTGATTTCATAGTTACCGGTGCCCAGC |

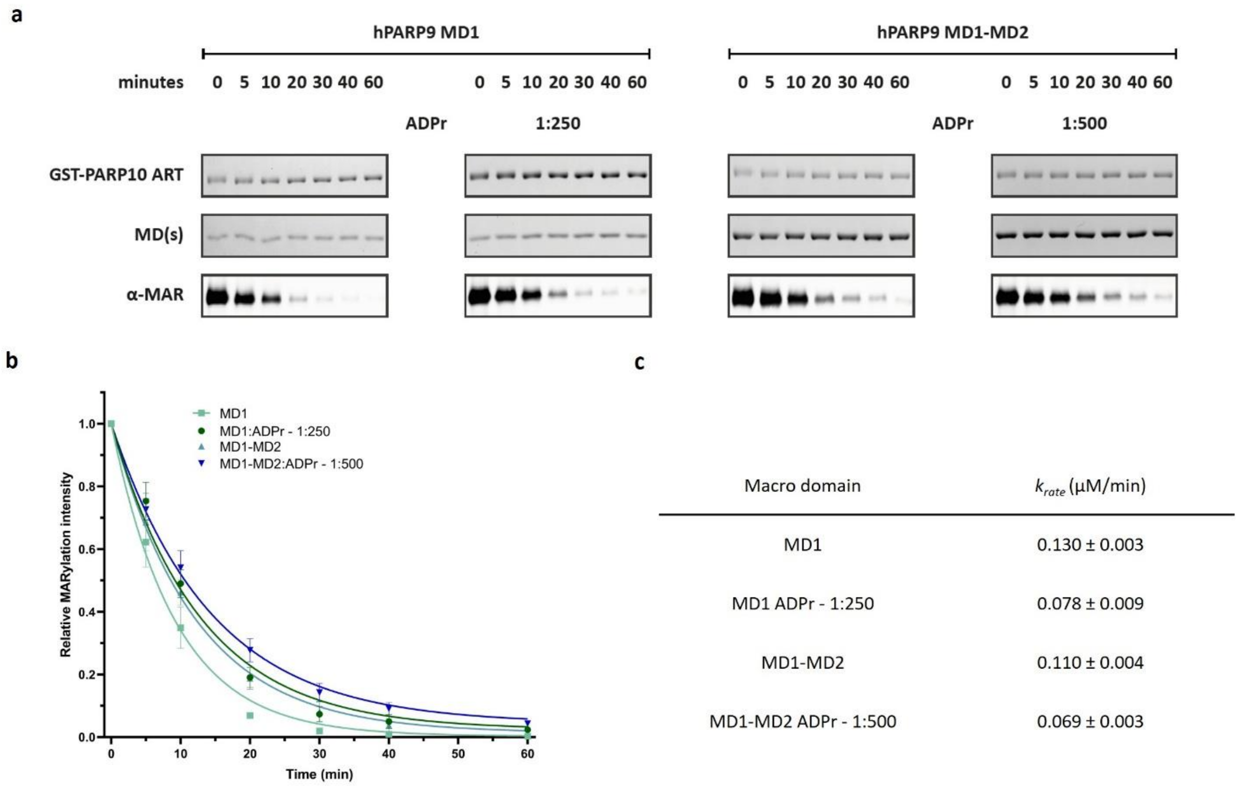


**Supplementary Figure 1. De-MARylation activity of hPARP9 MDs. a.** Time-course analysis of MARylated GST-PARP10 substrate de-MARylation with and without ADPr. Samples were resolved via Bis-Tris SDS-PAGE. Substrate MARylation was detected by Western blot using MABE1076 antibody (bottom panel). Protein loading for the GST-PARP10 substrate (top panel) and the hPARP9 macro domain constructs (middle panel) was confirmed by Coomassie Blue (CB) staining. **b.** Quantified de-MARylation kinetics. Relative intensities of the MABE1076 signal were normalized to t=0 and fit to a one-phase decay model. Data represent mean ± SD (n=3). **c.** Calculated apparent reaction rates (krate) derived from the non-linear regression fits in b. ADPr-bound samples were pre-incubated at the indicated protein:ligand molar ratios. Band intensities were quantified using Image Lab software (Bio-Rad), and kinetic plots and rate constants were generated using GraphPad Prism.

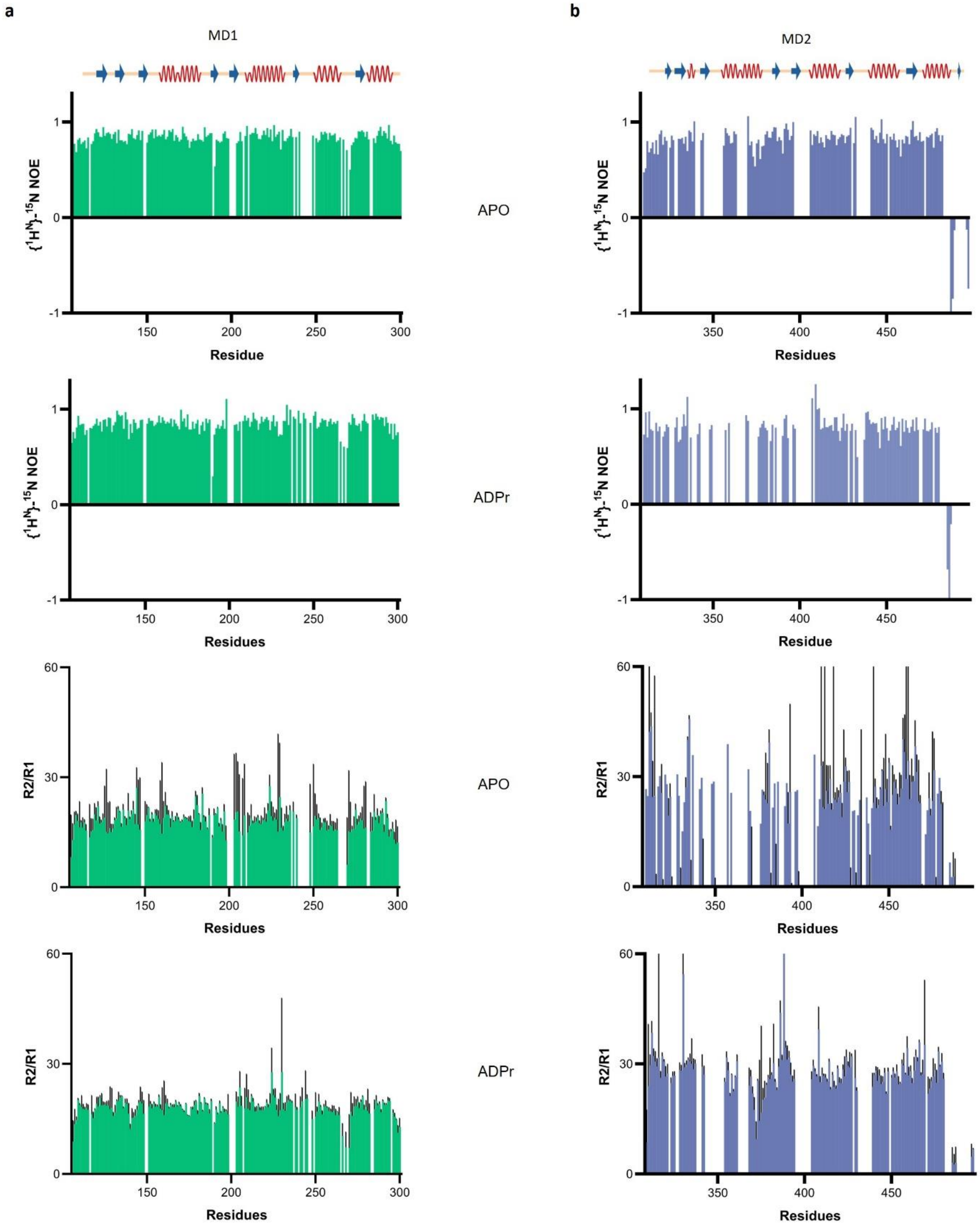
a
b
MD1
MD2
{$^1$H$^N$}-$^{15}$N NOE
R2/R1
APO
ADPr
APO
ADPr
Residue
Residues
Residues
Residue
1
0
-1
60
30
0
150
200
250
300
350
400
450

**Supplementary Figure 2. Backbone dynamics of PARP9 MD1 and MD2 macro domains.** Residue-specific $\{^1H\}$-$^{15}N$ heteronuclear NOE (top two rows) and $R_2/R_1$ ratios (bottom two rows) are plotted vertically for **a.** MD1 (green) and **b.** MD2 (purple) in their respective apo and ADPr-bound states. Secondary structure elements (β-strands as blue arrows, α-helices as red wavy lines) derived from crystallographic data are shown schematically above the profiles. NOE values tracking near 0.85 indicate a highly rigid, well-folded globular core on the ps-ns timescale, whereas significantly decreased or negative values (such as the C-terminus of MD2) highlight regions of elevated local fast-timescale flexibility. The $R_2/R_1$ ratios reflect global rotational diffusion, with the baseline levels mapping to the distinct rotational correlation times ($\tau_c$) of the domains. Pronounced $R_2/R_1$ spikes identify specific regions or residues undergoing conformational exchange ($R_{ex}$) on the μs-ms timescale. Gaps correspond to proline residues, unassigned backbones, or amide signals broadened beyond detection due to intermediate exchange within the binding pockets. All data were acquired at a proton frequency of 700 MHz and a temperature of 298 K.

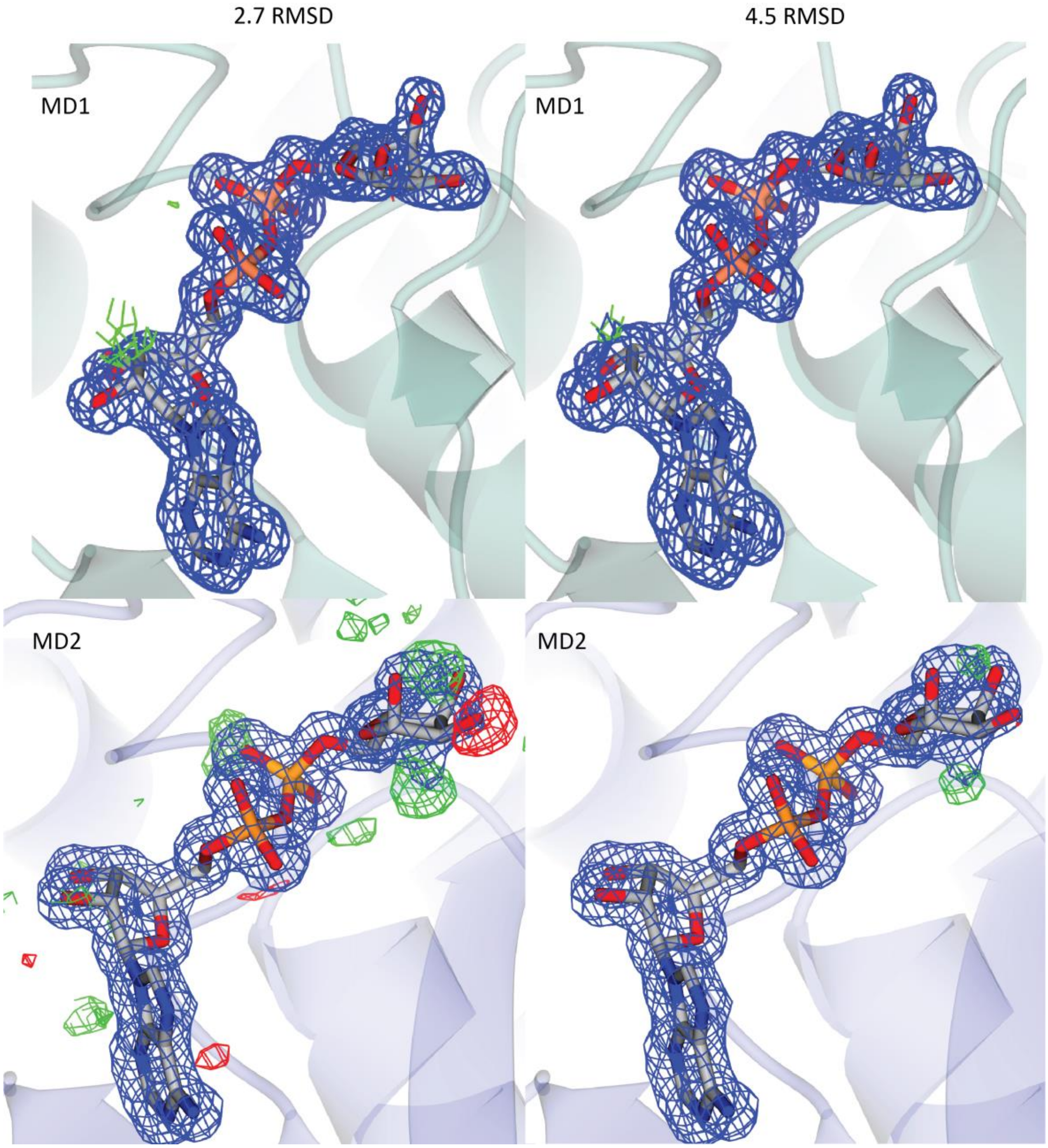


**Supplementary Figure 3. ADPr electron density in MD1 and MD2.** 2Fo-Fc electron density maps is contoured at 1.0σ. The difference electron density maps of ADPr in MD1 and MD2 binding pocket with only one anomeric form built are shown at 2.7 RMSD and 4.5 RMSD.

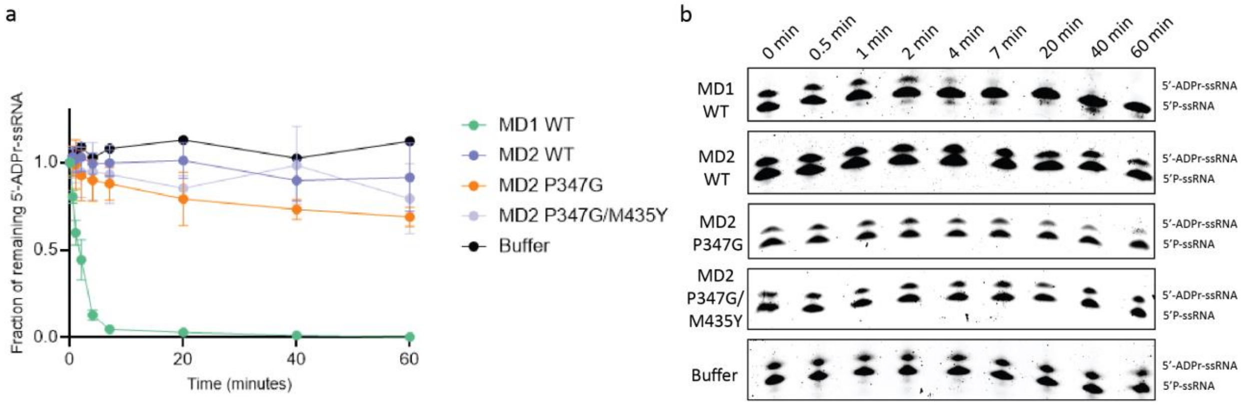


**Supplementary Figure 4. De-capping kinetics of PARP9 macro domains. a.** Time course of 5'-ADP-ribosylated ssRNA de-capping by MD1 WT, MD2 WT, MD2 P347G, MD2 P347G/M435Y, and buffer control. The substrate fraction was calculated as the ratio of substrate band intensity to total lane intensity (substrate + product) and normalized to t = 0. MD1 WT displays rapid, near-complete conversion, while MD2 P347G shows partial but reproducible activity. MD2 P347G/M435Y shows no detectable activity above the MD2 WT and buffer baselines, demonstrating that the M435Y substitution abolishes the gain-of-function conferred by P347G alone. Values represent mean ± SD (n = 3). The connecting lines are added for visualization purposes. **b.** Representative urea-PAGE gels showing conversion of the 5'-ADP-ribosylated ssRNA substrate (upper band) to the 5'-phosphate ssRNA product (lower band) over 60 minutes for each reaction variant. Band identities are indicated to the right of each gel.

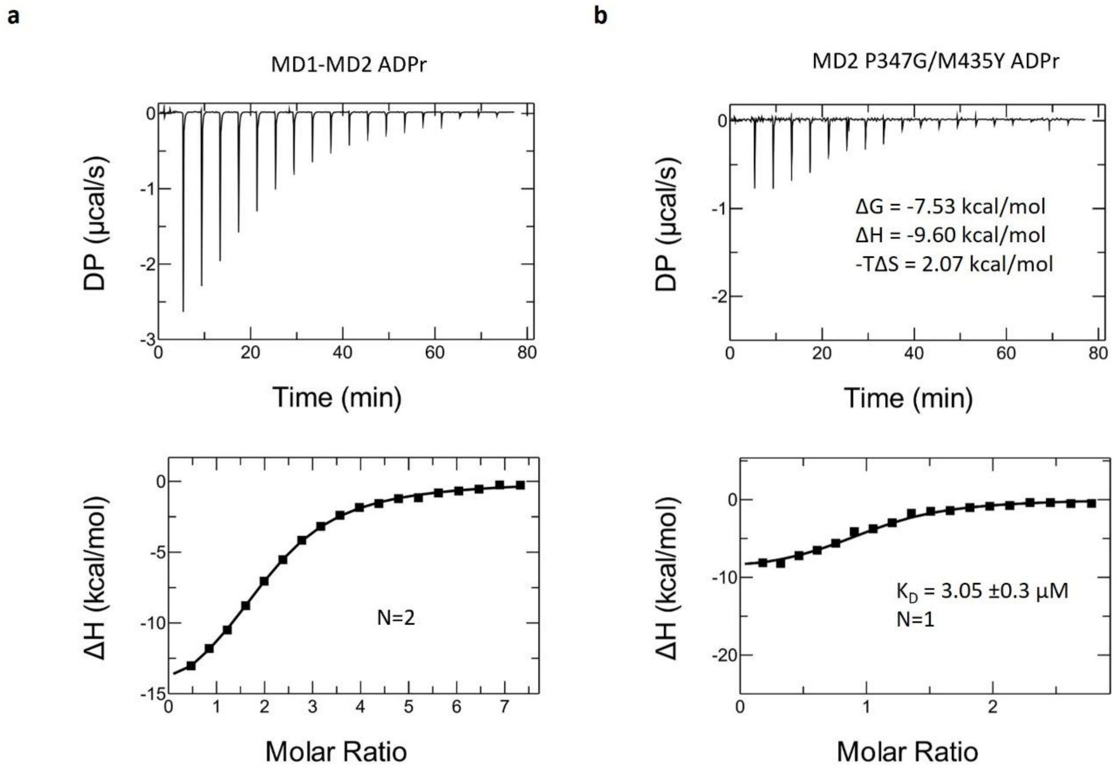


**Supplementary Figure 5. ITC analysis of ADP-ribose binding to the MD1-MD2 tandem construct and the MD2 P347G/M435Y double mutant. a.** ITC of ADP-ribose binding to the MD1-MD2 tandem. Upper panel: raw thermogram. Lower panel: integrated binding isotherm. The isotherm follows a single sigmoidal transition with N = 2, consistent with two ADPr binding events per tandem molecule. **b.** ITC of ADPr binding to the MD2 P347G/M435Y double mutant. Upper panel: raw thermogram with thermodynamic parameters. Lower panel: integrated binding isotherm fitted to a single-site model (N = 1).

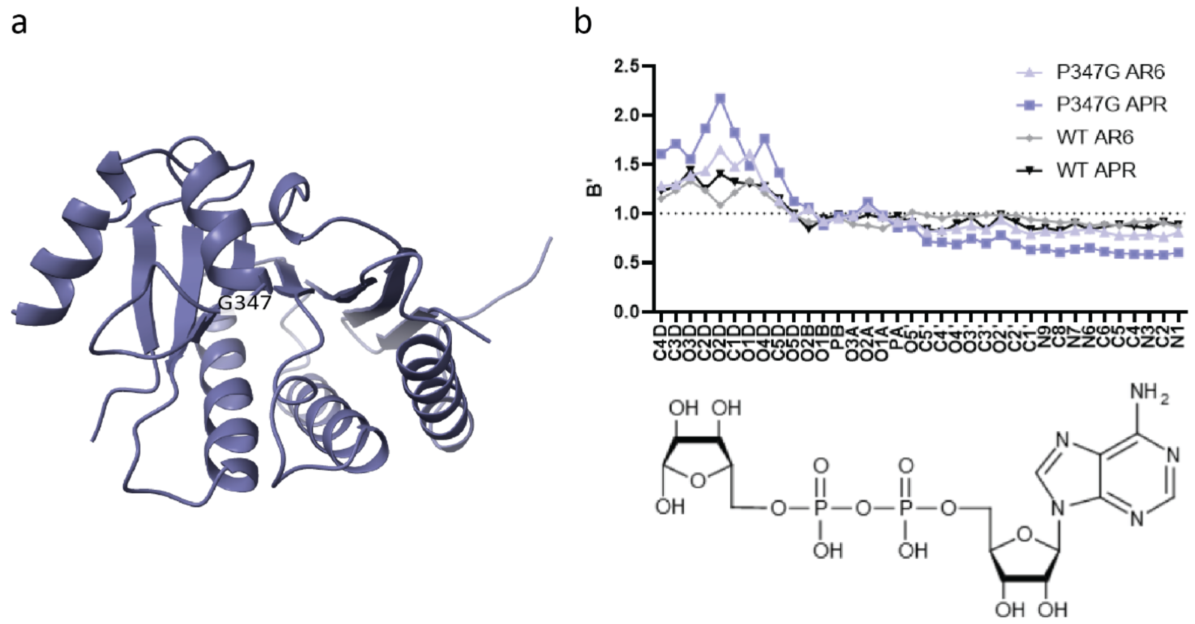


**Supplementary Figure 6. Structural analysis and ligand B-factor profiles of the PARP9 MD2 P347G mutant. a.** Ribbon representation of the crystal structure of MD2 P347G mutant in its apo state (32HB). The location of the mutated residue G347 is annotated. **b.** Normalized B-factor (B') profiles across individual atoms of the bound ligand for the α-anomer (AR6) and β-anomer (APR) of ADPr in WT and P347G MD2. A chemical schematic of ADPr is shown below for atom nomenclature reference.
